# How heterogeneous thymic output and homeostatic proliferation shape naive T cell receptor clone abundance distributions

**DOI:** 10.1101/674937

**Authors:** Renaud Dessalles, Maria R. D’Orsogna, Tom Chou

## Abstract

The set of T cells that express the same T cell receptor (TCR) sequence represent a T cell clone. The number of different naive T cell clones in an organism reflects the number of different T cell receptors (TCRs) arising from recombination of the V(D)J gene segments during T cell development in the thymus. TCR diversity and more specifically, the clone abundance distribution is an important factor in immune function. Specific recombination patterns occur more frequently than others while subsequent interactions between TCRs and self-antigens are known to trigger proliferation and sustain naive T cell survival. These processes are TCR-dependent, leading to clone-dependent thymic export and naive T cell proliferation rates. Using a mean-field approximation to the solution of a regulated birth-death-immigration model, we systematically quantify how TCR-dependent heterogeneities in immigration and proliferation rates affect the shape of clone abundance distributions (the number of different clones that are represented by a specific number of cells). By comparing predicted clone abundances derived from our heterogeneous birth-death-immigration model with experimentally sampled clone abundances, we quantify the heterogeneity necessary to generate the observed abundances. Our findings indicate that heterogeneity in proliferation rates is more likely the mechanism underlying the observed clone abundance distributions than heterogeneity in immigration rates.

**Author Summary:** The abundance distribution of different T cell receptors (TCRs) expressed on naive T cells depends on their rates of thymic output, homeostatic proliferation, and death. However, measured TCR count distributions do not match, even qualitatively, those predicted from a multiclone birth death-immigration process when constant birth, death, and immigration rates are used (a neutral model). We show how non-neutrality in the birth-death-immigration process, where naive T cells with different TCRs are produced and proliferate with a distribution of rates shape the predicted sampled clone abundance distributions (the clone counts). Using physiological parameters, we find that heterogeneity in proliferation rates, and not in thymic output rates, is the main determinant in generating the observed clone counts. These findings are consistent with proliferation-driven maintenance of the T cell population in humans.

## Introduction

Naive T cells play a fundamental role in the immune system’s response to pathogens, tumors, and other infectious agents. These cells are produced in the thymus, circulate through the blood, and migrate to the lymph nodes where they may be presented with different antigen proteins from various pathogens. Naive T cells mature in the thymus where the so-called V, D, and J segments of genes that code T cell receptors undergo rearrangement. Most T cell receptors (TCRs) are comprised of an alpha chain and a beta chain that are formed after VJ segment and VDJ segment recombination, respectively. The number of possible TCR gene sequences is extremely large, but while recombination is a nearly random process, not all TCRs are formed with the same probability. Before export to the periphery, T cells undergo a selection process, during which T cells with TCRs that react to self proteins are eliminated.

The unique receptors expressed on the cell surface of circulating TCRs enable them to recognize specific antigens; well known examples include the naive forms of helper T cells (CD4+) and cytotoxic T cells (CD8+). The set of naive T cells that express the same TCR are said to belong to the same T cell clone. Upon encountering the antigens that activate their TCRs, naive T cells turn into effector cells that assist in eliminating infected cells. Effector cells die after pathogen clearance, but some develop into memory T cells. Because of the large space of unknown pathogens, TCR clonal diversity is a key attribute in mounting an effective immune response. Recent studies also reveal that human TCR clonal diversity is implicated in healthy ageing, neonatal immunity, vaccination response and T cell reconstitution following haematopoietic stem cell transplantation [1, 2]. Despite the central role of the naive T cell pool in host defense, and broadly speaking in health and disease, TCR diversity is difficult to quantify. For example, the human body is known to host a large repertoire of T cell clones, however the actual distribution of clone sizes is not precisely known [3]. Only recently have experimental and theoretical efforts been devoted to understanding the mechanistic origins of TCR diversity [4–9]. The goal of this work is to formulate a realistic mathematical model that includes heterogeneous naive T cell generation and reproduction rates and that we will use to describe recent experimental results.

A well-established way to describe the T cell repertoire is by determining the clone abundance distribution or “clone count” 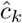 (for *k ≥* 1) that measures the number of distinct clones represented by exactly *k* T cells: 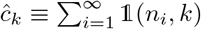, where *n*_*i*_ is the discrete number of T cells carrying TCR *i*. This distribution captures the entire pattern of the clonal populations. Several summary indices for T cell diversity such as Shannon’s entropy, Simpson’s index, or the whole population richness can be deduced from the distribution 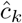[10]. Note that 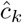 counts only the number of clones of a specific population *k* and does not carry any TCR sequence or identity information.

Complete clone counts 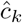 are difficult to measure in humans due to the large number of naive T cells (the total number of naive T cells is *N* ~ 10^11^ [11]). Nonetheless, high-throughput DNA sequencing on samples of peripheral blood containing T cells [12–15] have provided some insight into TCR diversity. A commonly observed feature is that all experimentally measured realizations of the clone counts 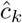 exhibit a power-law distribution in the clone abundance *k* [3, 4, 16–18]. Several authors have introduced stochastic models to explain the power law [4–7]. One of the proposed mechanisms is that T cells in different clones have TCRs that have different affinities for self-ligands that are necessary for peripheral proliferation [4–6], leading to clone specific replication rates. An alternative hypothesis [7] is that specific TCR sequences are more likely to arise in the V(D)J recombination process in the thymus [19] leading to a higher probability that these TCRs are produced. De Greef *et al.* [7] estimated the probability of production of a given TCR sequence by using the Inference and Generation of Repertoires (IGoR) simulation tool that quantitatively characterizes the statistics of receptor generation from both cDNA and gDNA data [19]. However, none of the previous studies have systematically incorporated heterogeneity in both immigration and replication rates, sampling, and comparison with measured TCR clone abundance distributions.

In this paper, we quantitatively analyze the effects of heterogeneity and sampling using a stochastic multiclone birth-death-immigration (BDI) process that allows for TCR-sequence dependent replication and immigration rates. Our model is based on a general continuous-time Markovian birth-death-immigration (BDI) process [10] where: (i) immigration represents the arrival of new clones from the thymus; (ii) birth describes homeostatic proliferation of naive T cells that yield newborn naive T cells with the same TCR as their parent; and (iii) death represents cell apoptosis. We also include a regulation, or “carrying capacity,” mechanism through a total population-dependent death rate which may represent the global competition for cytokines, such as Interleukine-7 [20–24], needed for naive T cell survival and homeostasis [25, 26]. This homeostasis will be considered as clone-independent since these cytokine signals are TCR-independent [22]. Mathematically, the inclusion of a carrying capacity ensures a finite naive T cell population at homeostasis.

We derive analytical results of our heterogeneous BDI model that are applicable on the scale of the entire organism. Our calculations provide insight into how parameters describing the shape of the distribution of immigration and proliferation rates affect the shape of the expected clone counts. To compare with experimental measurements, we also quantify the random sampling process that describes actual measurements derived from blood draws. Comparison with available TCR clone abundance data shows that predicted thymic output rate heterogeneity cannot generate the qualitative shape of the observed clone count distribution 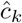. However, we show how a simple uniform proliferation rate distribution can yield the observed 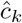.

## Materials and Methods

We focus on the clone counts 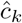 of the clone abundance distribution associated only with naive T cells, the first type of cells produced by the thymus that have not yet been activated by any antigen. Antigen-mediated activation initiates a one-way irreversible cascade of differentiation into effector and memory T cells that we can subsume into a death rate. Thus, we limit our analysis to birth, death, and immigration within the naive T cell compartment.

### Non-neutral Birth-Death-Immigration model

The multiclone BDI process is depicted in Fig. 1. We define *Q* to be the number of possible functional naive

**Fig 1.**
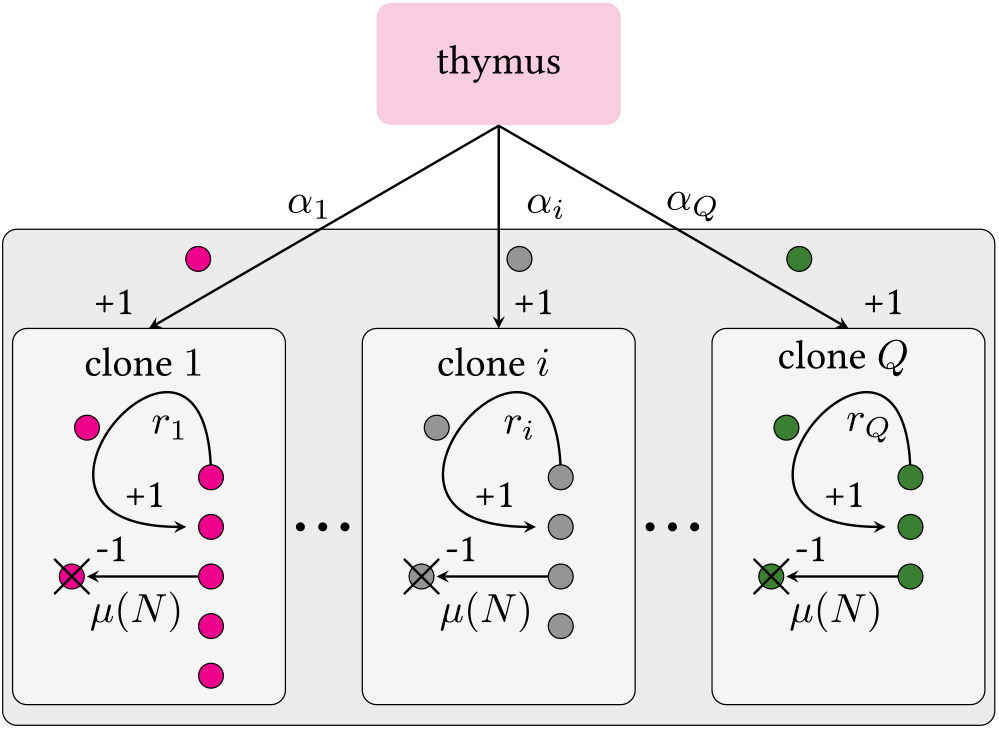
Schematic of a multiclone birth-death-immigration process. Clones are defined by distinct TCR sequences *i*. Theoretically, there are *Q* ≳ 10^15^ [6] or more [27, 28] possible viable V(D)J recombinations that lead to naive T cells. Due to finite and heterogeneous thymic output rates and and selection, the number of clones predicted in an entire organism is much less, on the order 10^6^ *−* 10^7^ [1, 6, 29–31]. Each clone carries its own thymic output and peripheral proliferation rates, *α*_*i*_ and *r*_*i*_, respectively. We assume all clones have the same population-dependent death rate *µ*(*N*). Since *Q* ≫ 1, we impose a continuous distribution over the rates *α* and *r*.

T cell clones that can be generated by V(D)J recombination in the thymus and that survive selection. This quantity is a small fraction (~ 1% [32]) of the total theoretical number of possible sequences, estimated to be 10^15^ − 10^18^ [6]. Thus, for each TCR chain, we choose *Q* = 10^15^ as the number of different clones that can be exported by the thymus. Due to naive T cell death, not all possible clone types will be present in the organism, so we denote the number of clones actually present in the body (or “richness”) by *C ≪ Q*, where estimates of *C* range from *∼* 10^6^ *−* 10^8^ in mice and human [1, 6, 29, 30]. The total number of naive T cells in an individual is *N ∼* 10^10^ *−* 10^11^ [11]. The discrete quantities *Q, C, N* are related to the discrete clone counts 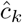 through

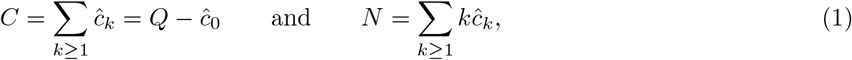

where 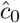 is the number of possible clones that are not expressed in the organism.

Each clone *i* (with 1 ≤ *i* ≤ *Q*) is characterized by an immigration rate *α*_*i*_ and a per cell replication rate *r*_*i*_. The immigration rate *α*_*i*_ is clone-specific because it depends on the preferential V(D)J recombination process; the replication rate *r*_*i*_ is also clone-specific due to the different interactions with self-peptides that trigger proliferation. Since the numbers of potential (*Q* ≫ 1) and observed (*C* ≫ 1) clones is extremely large, we can define a continuous density *π*(*α, r*) from which immigration and proliferation rates *α* and *r* are drawn. This means that the probability that a given clone has an immigration rate between *α* and *α* + d*α* and replication rate between *r* and *r* + d*r* is *π*(*α, r*)d*α*d*r*. Since *Q* is finite and countable, there are theoretical maximum values *A* and *R* for the immigration and proliferation rates, respectively, such that *π*(*α, r*) = 0 for *α* ≥ *A* and *r* ≥ *R*. In the BDI process, the upper bound *R* on the proliferation rate prevents unbounded numbers of naive T cells and is necessary for a self-consistent solution. Conversely, results are rather insensitive to the upper bound *A* on the immigration rate provided *π*(*α, r*) decays sufficiently fast such that the mean value of *α* is finite for all *r* ≤ *R*. Thus, for simplicity, we henceforth take the *A* → ∞ limit. The heterogeneity in the immigration and replication rates allow us to go beyond typical “neutral” BDI models, where both rates are fixed to a specific value for all clones.

Finally, we assume the per cell death rate *µ*(*N*) is clone-independent but a function of the total population *N*. This dependence represents the competition among all naive T cells for a common resource (such as cytokines), which effectively imposes a carrying capacity on the population [23, 28, 33]. We choose the linear form *µ*(*N*) = *µ*_1_*N* but the specific form of the regulation will not qualitatively affect our findings.

### Mean-Field Approximation

The exact steady-state probabilities over the discrete abundances 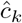 for a fully stochastic neutral BDI model with regulated death rate *µ*(*N*) were recently derived [10]. To incorporate TCR-dependent immigration and replication rates in a non-neutral model, we must consider distinct values of *α*_*i*_ and *r*_*i*_ for each clone *i*. In this case, an analytic solution for the probability distribution over 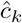, even at steady state, cannot be expressed in an explicit form. However, since the number of naive T cells (*N* ~ 10^11^ [11]) is large, we can exploit a mean-field approximation to the non-neutral BDI model and derive expressions for the mean values of the discrete clone counts 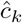. We will show later that under realistic parameter regimes, the mean-field approximation is quantitatively accurate. Breakdown of the mean field approximation has been carefully analyzed in other studies [34].

### Deterministic approximation for total population

To implement the mean-field approximation in the presence of a regulated death rate *µ*(*N*), we must first calculate the mean total steady-state population *N* ^∗^ ≫ 1. We start by writing the deterministic, “mass-action” ODE for the mean number *n*_*α,r*_(*t*) of cells with immigration rate *α* and proliferation rate *r* in a BDI process

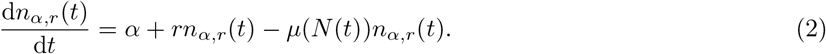

Since the number of clones that have an immigration rate between *α* and *α* + d*α* and a replication rate between *r* and *r* + d*r* is approximately *Qπ*(*α, r*)d*α*d*r*, the total mean number *N* (*t*) of naive T cells can be estimated as a weighted integral over all *n*_*α,r*_(*t*)

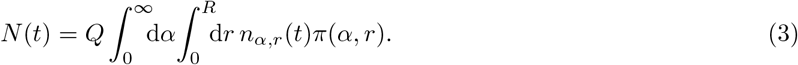

At steady-state, the solution to Eq. 2 can be simply expressed as

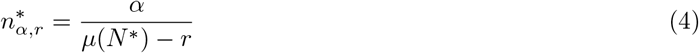

in which *N*^*∗*^ is the steady-state value of *N* (*t*). Thus, upon averaging Eq. 4 over *α* and *r*, we find

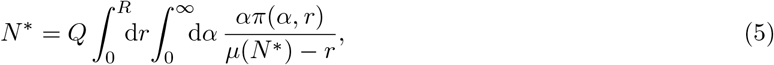

a self-consistent equation whose solution *N*^*∗*^ will be used in our mean-field approximation for the mean clone counts. Note that without the finite upper bound *R* of the density *π*(*α, r*), the integral in Eq. 5 diverges.

### Mean-field model of clone counts

We now use the results for the mean steady-state population *N*^*∗*^ to find the clone counts averaged over all realizations of the underlying stochastic process. The mean-field equations for the dynamics of these mean clone counts in the neutral model were derived in [34, 35] and are reviewed in Appendix A. In the neutral model, we assume that all effective clones *Q* are associated with the same rates *α* and *r* so that the mean field evolution equation for *c*_*k*_(*α, r*) is [34, 35]

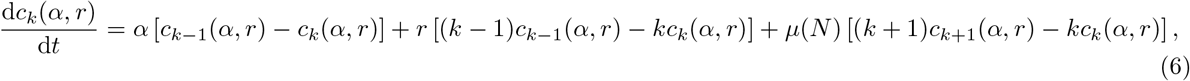

along with the constraint 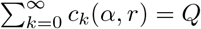. This equation assumes that both *c*_*k*_(*α, r*) and *N* are uncorrelated, allowing us to write the last term as a product of functions of the mean population 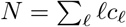 and *c*_*k*+1_, *c*_*k*_.

At steady state we replace *µ*(*N*) with *µ*(*N*\*), where the mean steady-state population *N*\* is found from self-consistently solving Eq. 5. The solution follows a negative binomial distribution with parameters *α/r* and *r/µ*(*N*\*) [10, 34]

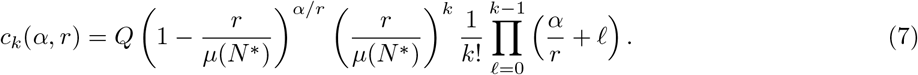

Now, suppose the clones obey possibly different values of *α* and *r*, as depicted in Fig. 1. The total mean clone count can be obtained from the *α, r*-specific result in Eq. 7 by averaging over *π*(*α, r*):

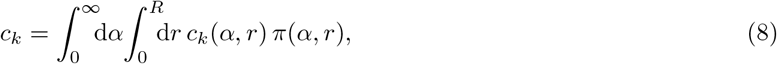

where for notational simplicity, we refer to the realization- and *π*(*α, r*)-averaged clones counts simply by *c*_*k*_. Note that since it is an average over the mean-field expression for *c*_*k*_(*α, r*), *c*_*k*_ defined in Eq. 8, is to be considered distinct from the actual 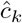, which is instead a discrete stochastic quantity that may be described via, *e.g.*, a *Q*-dimensional master equation. As mentioned earlier, the presence of heterogeneities makes computing the solution of the master equation impractical. Henceforth, we utilize the mean-field average *c*_*k*_ in Eq. 8 as a proxy for the discrete clone counts 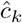, and discuss in detail the regimes where the identification 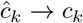 is justified; namely, cases where fluctuations may be considered small, and *c*_*k*_ can be compared to experimentally measured realizations. We also henceforth denote the mean field approximation to the total population and total richness by *N* and *C*, respectively. Since the population constraints are linear, these mean field quantities also obey the relationships in Eq. 1.

## Results

We now analyze Eqs. 5 and 8 using different choices for *π*(*α, r*) to show the effect of clone-specific immigration and proliferation rates on the overall measured clone abundance distribution. To determine the clone abundance *c*_*k*_ we must first solve the fixed point equation (Eq. 5) and use Eqs. 7 and 8 to determine *c*_*k*_. Rather than choosing a value for *µ*_1_ in *µ*(*N*) = *µ*_1_*N*, we first fix *N*\* to observed values and then determine *µ*_1_ that satisfies Eq. 5. The parameters used in our study are set according to

- The average total number of naive T cells is set to *N*^*∗*^ = 10^11^ [11].
- The total possible number of TCRs of either alpha or beta chains is set to *Q* = 10^15^ [36].
- The average immigration rate per clone 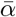 can be deduced from the total thymic output of all clones to be 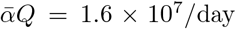 [37]. Using *Q* = 10^15^, the average per clone immigration rate is set to 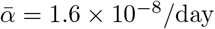.
- The average proliferation rate 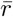 is estimated as 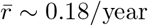. We thus set 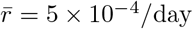 [37].

These parameters will be used in the rest of our analyses. We have verified that the predicted shapes of the clone counts are relatively insensitive to *Q* ≫ 1 and *N* * ≫ 1.

### Clone-specific immigration

To make analytical progress, we first assume that all clones have the same proliferation rate 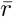 and that heterogeneity arises only in the immigration rate *α* through 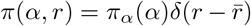. We analyze how differential V(D)J recombination probabilities affect the clone abundance distribution by drawing for each clone *i* (1 ≤ *i* ≤ *Q*), the immigration rate *α*_*i*_ from the probability density *π*_*α*_(*α*). By averaging the mean clone abundance (Eq. 8) over *π*_*α*_(*α*), we find

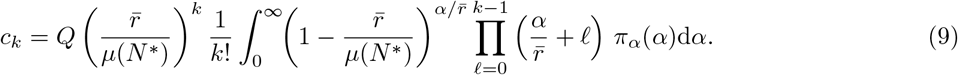

A specific form for *π*_*α*_(*α*) can be obtained from previous studies that predict V(D)J recombination frequencies associated with each TCR sequence. The statistical model for differential V(D)J recombination in humans is implemented in the Inference and Generation of Repertoires (IGoR) software [19]. In Appendix B.1, we estimate *π*_*α*_(*α*) by repeatedly running IGoR [19] and sampling a large number of TCRs. We assume that thymic selection is uncorrelated with V(D)J recombination so the relative probabilities of forming different TCRs provide an accurate representation of the ratios of the TCRs exported into the periphery. The observed frequency of each TCR shows that *π*_*α*_(*α*) can be approximated by a Pareto distribution with threshold parameter 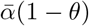 and shape parameter *θ*:

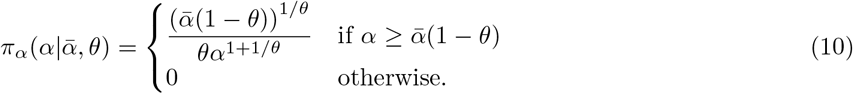

The parameters of the Pareto distribution will first be constrained such that 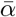 is set to the average per-clone immigration rate, 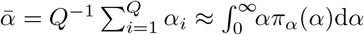.

The distributions *π*_*α*_(*α*) for different values of *θ* are shown in Fig. 2. An increase in the parameter *θ* will increase the variance of *π*_*α*_(*α*) (Fig. 2(a)-inset). In Fig. 2(b) we plot the mean clone count *c*_*k*_ derived in Eq. 9 with *π*_*α*_(*α*) specified by Eq. 10 for various values of *θ*. These curves are compared with those obtained when *θ* → 0 and 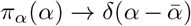 describing a fixed immigration rate 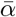. The BDI model in this neutral limit predicts a log-series distribution for *c*_*k*_ [10]. As can be seen in Fig. 2(b), deviations from the predictions of the neutral model (*θ* → 0) are noticeable only when *θ* ≳ 0.35. The estimates using IGoR fixes *θ* ≈ 0.28, suggesting that heterogeneity in thymic output is insufficient for producing the observed clone abundance distributions of naive T cells.

**Fig 2.**
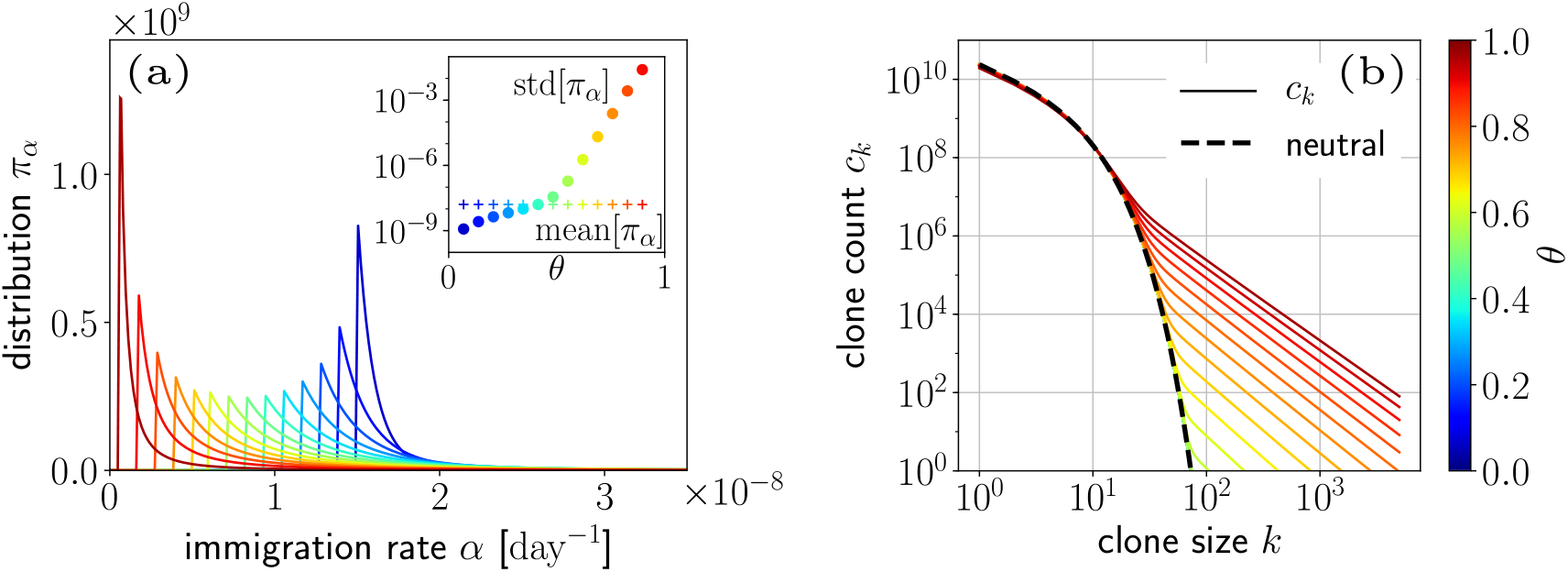
Effects of immigration rate heterogeneity. (a) Different Pareto distributions 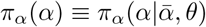 of the immigration rate *α* plotted for the same value of the mean 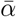 but different values of the shape parameter *θ* in Eq. 10. Increasing *θ* increases the heterogeneity in the immigration rate per clone *α*. Although the different distributions appear to have different mean values, the longer tails of the distributions with larger *θ* yield means that are invariant to *θ*. Inset: Standard deviation of *π*_*α*_(*α*) as a function of *θ*. (b) Expected clone counts *c*_*k*_ for different choices of *θ* (Eq. 9). The clone counts differ significantly from that of the BDI neutral model only when *θ* ≳ 0.35, as can be seen from the dashed curve plotted above.

The shape of *c*_*k*_ as derived from Eqs. 9 and 10 exhibits two regimes. At small clone sizes *k*, regardless of the value of *θ*, *c*_*k*_ follows that of the neutral model. At larger sizes *k*, *c*_*k*_ converges to a power-law. The transition between these two regimes occurs at a characteristic size that decreases with *θ*, and with the standard deviation of *π*_*α*_(*α*). In Appendix B.2 and Fig. S2(b), we show that clones within these two regimes correspond to two sub-populations. The first, with smaller clone sizes, is characterized by immigration rates *α* smaller than the proliferation rate 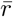. For these clones, the production of new cells is due mostly to peripheral proliferation rather than from thymic immigration. The second sub-population with larger clone sizes corresponds to clones with immigration rates 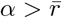. The production of new cells in these clones is driven mainly by thymic immigration rather than by proliferation. Intuitively, one can see that as *θ* increases, the proportion of clones with higher immigration rate *α* increases as well, leading to a larger representation of the second, high immigration rate sub-population.

### Clone-specific proliferation

The other limit of clonal heterogeneity that can be readily analyzed is that of an immigration rate that is common to all clones and proliferation rates that are clone-dependent. The peripheral proliferation of T cells depends on the affinity of the corresponding TCR to self-antigens. TCR-antigen affinity in turn depends on the receptor amino acid sequence; thus, the rate at which a T cell proliferates can be clone-specific [28, 38]. The distribution of proliferation rates among all the *Q* possible clones is a mapping to the interactions between TCRs and low-affinity MHC/self-peptide complexes that has not been experimentally resolved.

For simplicity, we assume 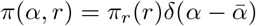 where 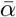 is the common immigration rate and where the probability that a clone has a replication rate between *r* and *r* + *dr* is *π*_*r*_(*r*)d*r*. We previously introduced the biologically relevant constraint that *π*(*r*) = 0 for *r > R*; we now further assume that *R < µ*(*N*\*) so that a finite fixed point solution *N*\* exists for Eq. 5. This assumption implies that the death rate is larger than the fastest proliferation rate and allows us to write

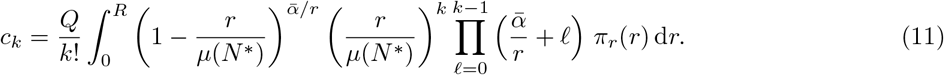

We illustrate in Fig. 3 the effect of a uniform proliferation rate distribution

**Fig 3.**
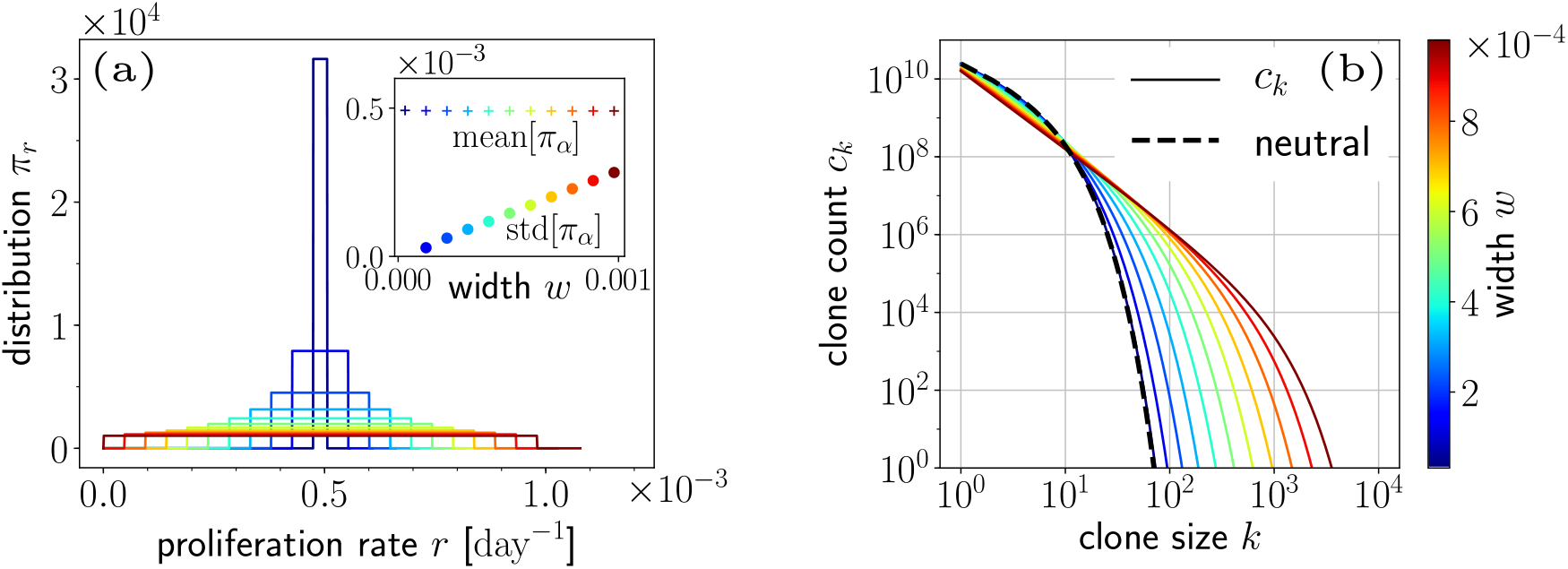
Effects of proliferation rate heterogeneity. (a) Different “box” distributions for *π*_*r*_(*r*) associated with different widths *w* (see Eq. 12). Larger *w* corresponds to higher heterogeneity across clones in the per-cell proliferation rate *r*. Inset: Standard deviation of *π*_*r*_(*r*) for different values of *w*. (b) Predicted clone abundances *c*_*k*_ from Eq. 11 for different choices of *w*. Wider distributions generate longer-tailed distributions.

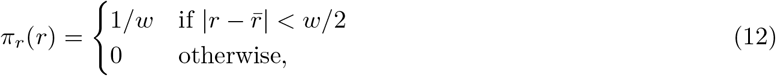

where *w* represents the width of the uniform distribution centered about 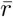, as shown in Fig. 3(a). By construction, the maximum allowable immigration rate is 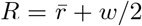. Fig. 3(b) shows that an increase in *w* leads to an increase in the proportion of large clones. It is important to note that the shape of the clone abundance *c*_*k*_ is particularly sensitive to the behavior of *π*_*r*_(*r*) near the upper bound *R* as can be seen in Fig. S3. This is because, regardless of their initial numbers, clones with *r ≈ R* will have a proliferation advantage over other clones and will eventually dominate the total naive T cell population, as shown in Fig. S3(c).

### Sampling

Unless an animal is sacked and its entire naive T cell population sequenced, TCR clone distributions are typically measured from sequencing TCRs in a small blood sample. In such samples, low population clones may be missed. In order to compare our predictions with measured clone abundance distributions, we must revise our predictions to allow for random cell sampling.

We define *η* as the fraction of naive T cells in an organism that is drawn in a sample and assume that all naive T cells in the organism have the same probability *η* of being sampled. This is true only if naive T cells carrying different TCRs are not preferentially partitioned into different tissues and are uniformly distributed within an animal. Let us assume that a specific TCR is represented by ℓ cells in an organism. If *N*_*η*_ ≫ ℓ, the probability that *k* cells from the same clone are sampled approximately follows a binomial distribution with parameters ℓ and *η* [39–41]

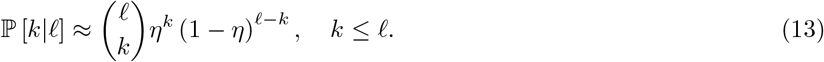

The associated mean *sampled* clone count 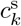 depends on the clone count *c*_ℓ_ in the body via the formula

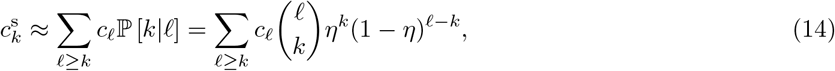

where *c*_ℓ_ is found through Eq. 8.

In Fig. 4(a) we show the effects of sampling by plotting both the whole-organism *c*_*k*_ derived from Eq. 8 and the corresponding sampled distribution 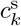 calculated using Eq. 14. We show three cases: i) the *c*_ℓ_ distribution is derived from a fully neutral model with specific values of 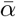 and 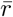; ii) the proliferation rate is fixed at 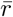 and a high heterogeneity in the immigration rate *α* is chosen with *π*_*α*_(*α*) given in Eq. 10 with *θ* ~ 1, iii) the immigration rate is kept fixed at 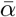 and proliferation rates *r* are highly heterogeneous with 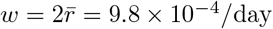 in *π*_*r*_(*r*), as given in Eq. 12.

**Fig 4.**
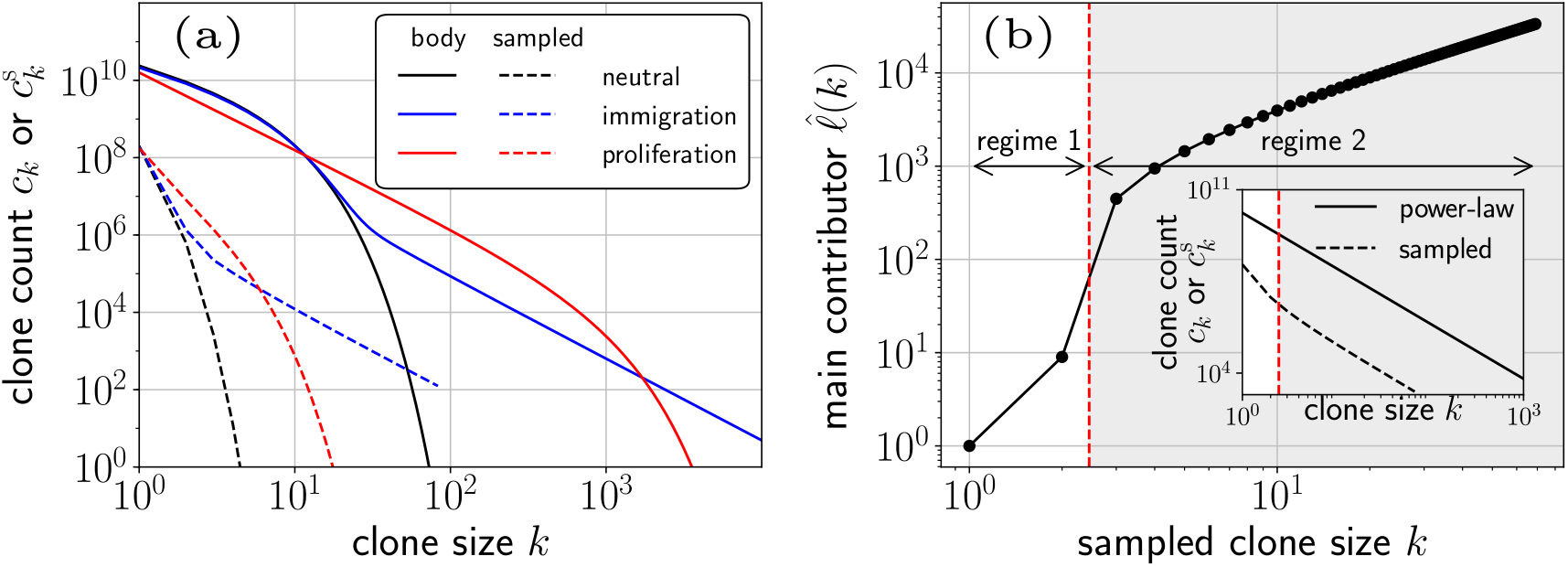
Effect of sampling on mean clone abundances *c*_*k*_. (a) Whole-body mean clone abundances *c*_*k*_ (solid curves) and expected sampled (dashed curves) abundances 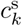 for the neutral (black), immigration rate heterogeneity (blue), and proliferation rate heterogeneity (red) models. The sampling fraction used is *η* = 2 × 10^*−*3^. (b) Illustration of two regimes in sampling. We considered sampling from an idealized pure power-law clone abundance *c*_ℓ_ ~ ℓ^−λ^, λ = 2.1 (inset) that is consistent with that predicted by the heterogeneous proliferation rate model (solid red curve in (a)). For a sampled clone of size *k*, the main plot shows the most likely abundance of said clone in the whole body, 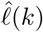, defined in Eq. 15. Plotting 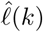 indicates two regimes in the sampling. The main contributors to sampled clones of size *k* ≤ 2 (regime 1) are smaller whole-body clones (ℓ≲ 10), while the main contributors to clones of size *k* ≥ 3 (regime 2) are clones of size ℓ ≳ 10^3^.

Experimental sampling will strongly affect the observed clone abundances 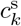 especially when *η ≪* 1. We set the sampling fraction to *η* = 2 *×* 10^*−*3^, which is typical for data on humans. In all three cases described above the observed clone abundances 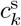 are significantly reduced with respect to *c*_*k*_, reducing the overall slopes of the power-law-like clone abundance distributions. The magnitude of the reduction in the exponent depends on the value of *η* and can be quite significant as can be seen in Fig. S4).

An interesting crossover feature can be seen, particularly when sampling in the case of heterogeneous immigration rates, as described in case ii) above. As can be seen from the blue curves in Fig. 4(a), two power-law regimes arise with a crossover size *k* ~ 2 − 3. To understand this behavior, consider the effect of sampling from a hypothetical power-law clone abundance *c*_ℓ_ ~ ℓ^−λ^, λ = 2.1 as shown in the inset of Fig. 4(b). For a given sampled clone of size *k*, we can calculate its most probable size 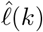 in the organism by evaluating the maximum likelihood of the binomial sampling

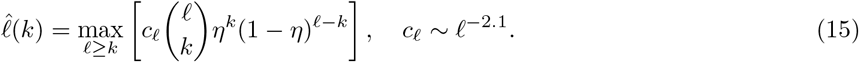

The 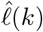 curve in this case is shown in Fig. 4(b). For clones of size *k* ≤ 2 measured in the sample, the most likely size in the organism 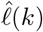 is also small, less than ~10 copies in the body. Conversely, sampled clones of size *k* ≥ 2 are most likely to have originated from clones with ≳ 10^3^ copies in the body. The two regimes observed in Fig. 4(b) can then be interpreted as a trade-off between the contributions of the large number of clones with small populations and the few number of clones with large populations, leading to a cross-over behavior in *c*_*k*_. This cross-over arises in the heterogeneous immigration model (blue curve) in Fig. 4(a) and is a consequence of sampling and not the intrinsic kink in the whole-body *c*_*k*_ at *k* ≈ 30. The sampling trade-off is not apparent in the neutral and heterogeneous proliferation model, case i) and iii) above, and depicted by the black and red curves in Fig. 4(a). These distributions decay too fast before the “sampling kink” reveals itself.

### Comparison with measured clone abundances

We now compare our predictions for 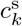 with measured clone abundances from Oakes *et al.* [13]. These data were extracted by combining barcoding with TCR sequencing of either the alpha or beta chains of TCRs from naive human CD4+ and CD8+ T cells, thereby eliminating PCR bias, especially for larger clones [15]. In Fig. 5 we plot the experimental clone abundances alongside our theoretical results for 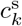 as evaluated in Eq. 13. We focused our fitting to small- and intermediate-sized clones since the mean field approximation at large clone sizes *k* is not accurate [34], the frequencies at large *k* are low and fitting to a continuous expected distribution is not reliable, and the single large clones represented by the flat region in Fig. 5 contain memory T cells that have expanded [26, 42]. Moreover, we do not expect a model for an expected continuous clone count to accurately capture these isolated, discrete single clones (open circles) in the large-*k* regime where *c*_*k*_ = 0, 1. Since our model does not incorporate the emergence of memory T cells, and the large-*k* data is more likely to include a higher percentage of memory cells, we simply drop the single-clone counts and fit only those alpha chain or beta chain clones for which the clone count *c_k_ ≥* 2.

**Fig 5.**
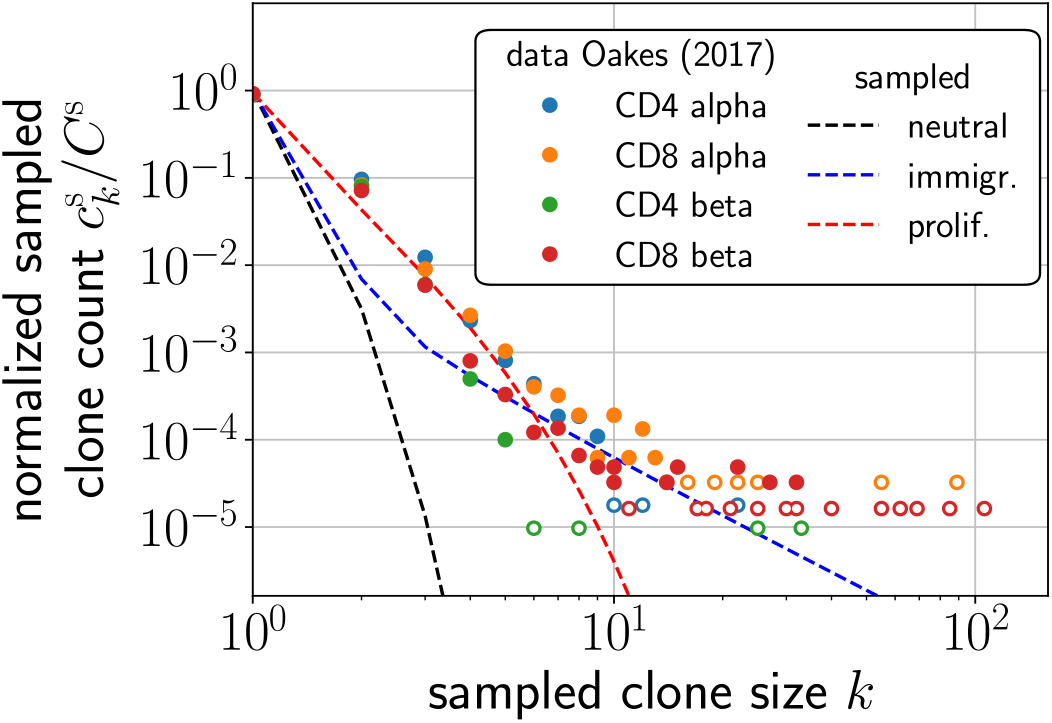
Comparison of the predicted mean sampled clone abundances 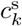 (dashed) with measured human TCR alpha and beta chain seuence abundances (dots) in which PCR bias was avoided [13]. The best-fit model (for reliable measurements at modest sizes *k*) model includes proliferation rate heterogeneity. In the dataset shown, TCRs observed in memory cells are not subtracted. Data points corresponding to single clones are denoted by the open circles.

As shown by the black curve in Fig. 5, the expected sampled clone count 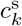 derived from the neutral model with fixed 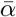, 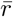 cannot qualitatively fit the data. Heterogeneity in immigration rate *α* also does not yield a good match to data for realistic values of the sampling parameter *η*, except possibly at large clone sizes *k* where the clone counts are highly variable, and only for high degrees of heterogeneity *θ* ~ 0.9. However, as estimated from IGoR, *θ* ≈ 0.28 *>* 0.9. At this value of *θ*, no significant difference between the neutral and the heterogeneous immigration rate cases are to be expected, especially after sampling. Thus, a heterogeneous immigration rate model cannot be consistent with both measured clone abundances [13] and the level immigration heterogeneity estimated from IGoR [19].

On the other hand, the red curves in Fig. 5 show that the 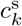 predicted using heterogeneity in the proliferation rate match the data for reasonable values of the clone size *k*, using realistic sampling fractions *η*. In this context, since a specific TCR is composed of one alpha chain and one beta chain, and since the distributions of alpha and beta chains are quantified separately, the heterogeneous proliferation rates should be interpreted as those of an individual alpha or beta sequence, and not as the heterogeneity in functional TCR-self-antigen avidity.

In Fig. S4, we show additional comparisons for different values of the sampling fractions, confirming that *η* ~ 2 × 10^*−*3^ provides the optimal fitting with measured clone counts. In summary, it is evident that the neutral model cannot be made to fit the data without using unrealistic sampling rates *η* ≳ 0.1. The model with large immigration heterogeneity also fails to fit some of the data, in particular, those derived from sequencing different beta chains on CD4+ cells.

We have also tried to fit our model to data from mice, such as from Zarnitsyna *et al.* [31]. These preliminary data did not filter out PCR errors and yielded biphasic clone abundance distributions with *λ* ≈ 1.76, 1.1. The magnitudes of these exponents are too small to be accurately fit by a heterogeneous BDI process, even when unrealistic distributions in *r* are used and the effects of sampling are neglected.

## Discussion and Conclusions

Here, we review and justify a number of critical biological assumptions and mathematical approximations used in our analysis. The effects of relaxing our approximations are also discussed.

### Mean field approximation

Our mean-field approximation is embodied in Eq. 6, where correlations between fluctuations in the total population 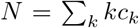 in the regulation term *µ*(*N*) and the explicit *c*_*k*_ terms are neglected. This approximation has been shown to be accurate for *k* ≲ *N*\* when 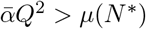 [34]. When the total T cell immigration rate is extremely small, the above constraint is violated and a single large clone arises due to competitive exclusion [34, 43, 44]. In this case, an accurate approximation for the steady-state clone abundance *c*_*k*_ can be obtained using a variation of the two-species Moran model [34]. Thus, except under such extreme scenarios, the parameters associated with the human adaptive immune system satisfy the condition for the mean-field approximation to be accurate.

For large *k* the number of cells contributing to *c*_*k*_ is also large so demographic stochasticity is relatively small and results in small uncertainties in the value of *k*, and not in the magnitude of *c*_*k*_. However, the mean field approximation to large-*k* clone counts is not accurate. For very large clone sizes, the total population constraint forces *c*_*k*_ to rapidly approach zero for *k* ≳ *N*\*, a feature that is not accurately reflected in the mean-field approximation [34]. Finally, large clones likely include memory T cells that have been produced after antigen stimulation of specific clones. Thus, the data at large *k* is likely to contain T cells in the memory pool. Thus, in our analysis, we perform qualitative “fits” to the data only for modest values of *k*.

### Estimation of *π*(*α, r*)

Given an explicit mean-field solution to *c*_*k*_(*α, r*) from Eq. 7, distributions *π*(*α, r*) detailing the immigration and proliferation rates *α* and *r* are then required to compute the expected clone abundances in *c*_*k*_ from Eq. 8. The main difficulty is accurately representing immigration and proliferation heterogeneity. Even if we assume that the immigration and proliferation rates are uncorrelated and that *π*(*α, r*) is factorizable, *π*(*α, r*) = *π*_*α*_(*α*)*π*_*r*_(*r*), there have been no direct experimental measurements of either *π*_*α*_(*α*) or *π*_*r*_(*r*).

We refer to a study on the statistical inference of the probability of TCR sequences to estimate immigration heterogeneity. In particular, we used the IGoR (Inference and Generation of Repertoires) computational tool which predicts recombination events [19]. We selected approximately 10^8^ such IGoR recombination events, and created a frequency histogram on the observed sequences. The resulting frequency distribution closely follows Zipf’s law [45, 46], from which we constructed a self-consistent Pareto distribution for *π*_*α*_(*α*), with a mean 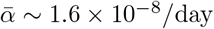 [37]. As mentioned, we assume that selection and recombination are uncorrelated and that the results from IGoR can be used to infer the heterogeneity of the thymic output after selection. While IGoR allows us to reconstruct *π*_*α*_(*α*) there are no reliable data on the *in vivo* distribution of TCR-dependent homeostatic proliferation rates *r*.

Previous studies modeled the proliferation rate of each TCR as being directly proportional to the average interaction times between the TCR itself and a large number of self-ligands [4, 6]. Each TCR was assumed to interact with multiple self-antigens through cross-reactivity and the interaction of each TCR–self-antigen pair is sampled from a given Gaussian [4] or log-normal [6] distribution. However averages were taken over a large number of TCR–self-ligand interactions sampled from the same distribution and may underestimate the true proliferation heterogeneity. In this work we assumed a simple box distribution for *π*_*r*_(*r*) centered around the mean value 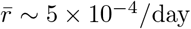 [37] and analyzed the predicted the clone abundances as the width *w* of the box is varied.

### Qualitative description of clone abundances *c*_*k*_

Here, we offer a qualitative interpretation of the observed clone abundance profiles *c*_*k*_ as a function of *k*, as seen in Fig. 4. The arguments we use closely resemble those presented to describe the two regimes observed under heterogeneous immigration, case ii), and are more thoroughly discussed in Appendix B.2 and Fig. S2(b). We begin by dissecting the behavior of *c*_*k*_(*α, r*), which, as described above, defines a negative binomial distribution with parameters *u* = *α/r* and *v* = *r/µ*(*N*\*). In Fig. 6 we plot the mean clone size *k*,

**Fig 6.**
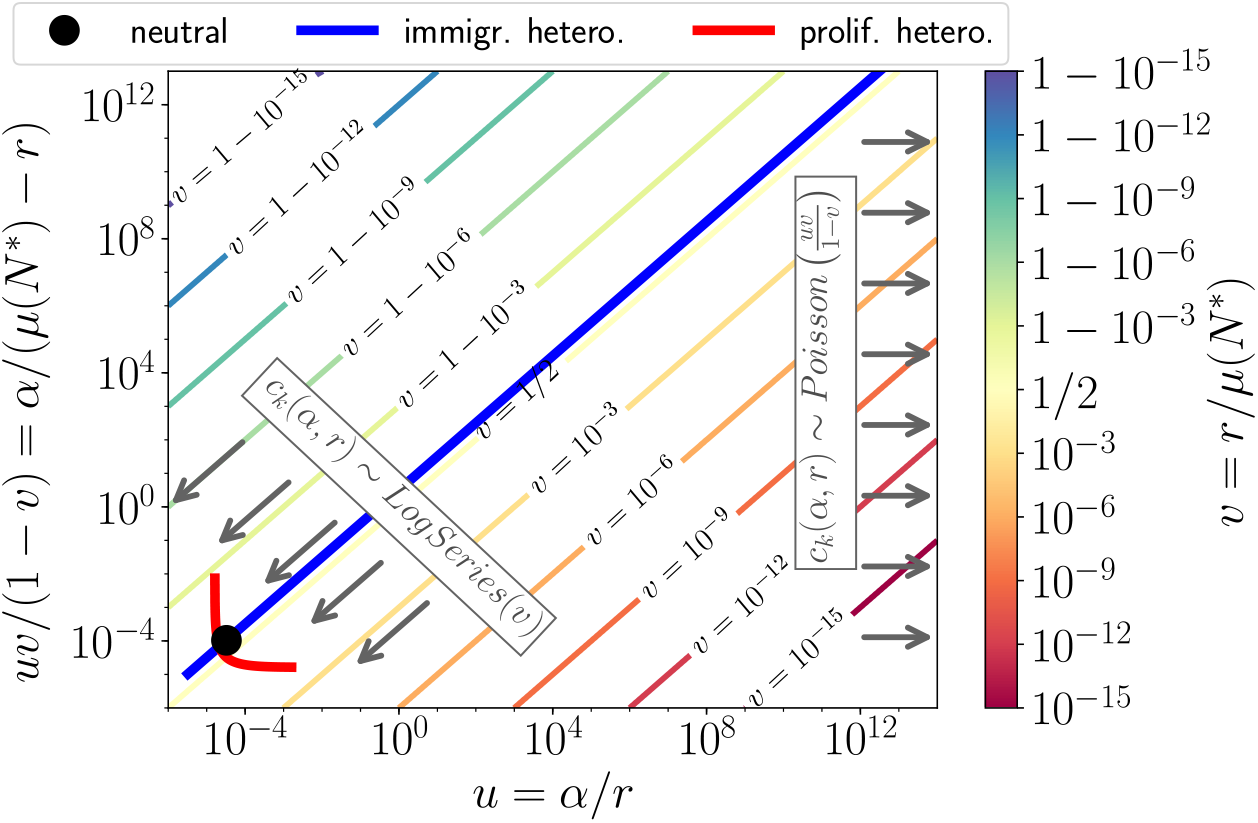
Parameter regimes of different distributions. For a clone with immigration rate *α* and proliferation rate *r*, (whose clone size follows a negative binomial distribution of parameters *u* = *α/r* and *v* = *r/µ*(*N*\*), see Eq. 7, the diagram shows the average clone size *uv/*(1 − *v*) = *α/*(*µ*(*N*\*) − *r*) as a function of *u* = *α/r*. When *v >* 0 is held constant and *u* → 0, the clone counts (conditioned to *k >* 0) converges to a log series distribution. If *uv/*(1 − *v*) = *α/*(*µ*(*N*\*) − *r*) remains constant while *u* → ∞, the clone size converges to a Poisson distribution. The “line integral” of the three models of Fig. 4 are shown: both the neutral model, and the proliferation model are in the log-series regime; the immigration regime covers both the log-series and the Poisson regime.

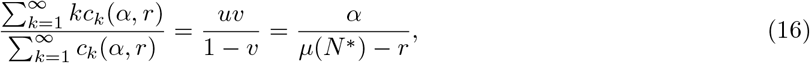

as a function of *u* = *α/r*. Two different regimes emerge:

- If *u* → 0 while *v* = *r/µ*(*N*\*) remains constant (lower left in Fig. 6), *c*_*k*_(*α, r*) (conditioned on *k >* 0) converges to a log-series distribution.
- If *u → ∞* while the averaged clone size given by *uv/*(1 *− v*) = *α/*(*µ*(*N*\*) *− r*) (Eq. 16) remains constant, *c*_*k*_(*α, r*) converges to a Poisson distribution with parameter *uv/*(1 *− v*).

These results follow from analogous considerations made in Appendix B.2. Finally, we show paths in the *u* = *α/r* and *uv/*(1 − *v*) = *α/*(*µ*(*N*\*) − *r*) plane over which the constrained “line integrals” correspond to the distributions *π*(*α, r*) for the neutral, heterogeneous immigration, and heterogeneous proliferation rate models.

Results are displayed in Fig. 6. As can be seen, both the heterogeneous proliferation and the neutral cases fall in the log-series regimes; here the expected clone counts result from an integration of different log-series distributions over *π*(*α, r*). The heterogeneous immigration case includes both log-series and Poisson distributions, generating the two qualitatively different regimes. More details are discussed in Appendix B.2).

### Correlated immigration *α* and proliferation *r*

Hitherto, we have considered independent immigration and proliferation, and assumed a factorizable rate distribution *π*(*α, r*) = *π*_*α*_(*α*)*π*_*r*_(*r*). However, immigration and proliferation rates may be correlated for certain clones. For example, a frequent realization of V(D)J recombination may also result in a TCR that is more likely to be activated for proliferation. In this case, *α* would be positively correlated with *r*. In Fig. S5 we consider the effect of correlated *π*(*α, r*). For 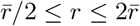, we considered normalized, positively/negatively correlated box distributions as shown in Fig S5(a):

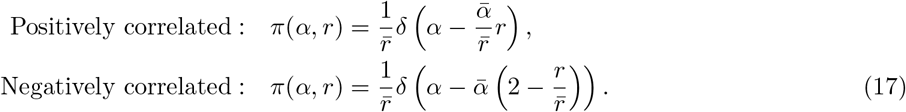

Within our mean field model, these correlated distributions *π*(*α, r*) result in very similar expected clone abundance distributions *c*_*k*_ (Fig S5(b)). This insensitivity to correlations between immigration and proliferation can be qualitatively understood by considering the “line integral” over dominant paths of *π*(*α, r*) in the *uv/*(1 − *v*) = *α/*(*µ*(*N*\*) − *r*) *vs. u* = *α/r* diagram. As shown in Fig. S5(c), both line integrals remain in the log-series distribution regime, indicating that the clone abundance distributions are qualitatively similar to that predicted by a model with proliferation heterogeneity alone.

### Steady state assumption

In this study, we only considered the steady state of our birth-death-immigration model in Eq. 7 because this limit allowed relatively easy derivations of analytical results. This was also the strategy for previous modeling work [4, 6, 7, 34, 35]. However, the per clone immigration and proliferation times may be on the order of months or years, a time scale over which thymic output can change significantly as an individual ages 33, 37, 47, 48] and as thymic output diminishes [33, 37]. Moreover, clone abundance distributions have been shown to show specific patterns as a function of age [49–51].

From the dynamics of *N* (*t*) in a neutral model with fixed 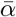 and 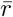, the timescale for relaxation to steady state can be estimated to be *∼* 15 *−* 20 years [33]. Besides the total population, the different subpopulations of specific sizes described by their number *c*_*k*_ relax to steady-state in a spectrum of time scales depending on the clone sizes *k* [52].

Thus, the steady-state may not be strictly reached and should be considered as an approximation, especially given time-dependent perturbations, including aging, to the adaptive immune system of an individual.

These timescales can be approximated by the eigenvalues of linearized forms of Eqs. 6. Besides time-dependent changes in *α*, more subtle time-inhomogeneities such as changes in proliferation and and death rates have been demonstrated [47, 48]. Thus, our steady-state assumption should be relaxed by incorporation of time-dependent perturbations to the model parameters *µ*(*N*) and/or *π*(*α, r*). Longitudinal measurements of clone abundances or experiments involving time-dependent perturbations would provide significant insight into the overall dynamics of clone abundances.

### General conclusions

We developed a heterogeneous multispecies birth-death-immigration model and analyzed it in the context of T cell clonal heterogeneity; the clone abundance distribution is derived in the mean-field limit. Unlike previous studies [4], our modeling approach incorporated sampling statistics and provided simple formulae, allowing us to predict clone abundances under different rate distributions for arbitrarily large systems (*N ∼* 10^11^), without the need for simulation.

The heterogeneous BDI model produces mean clone count distributions that follow power laws over a range of sizes *k*, with varying exponents that depend on the immigration and proliferation rate heterogeneity and sampling size *η*. We were able to compare results from our model to measured clone abundance distributions in alpha and beta TCR chains. Based on recombination statistics inferred from cDNA and gDNA sequences [19], we derived a Pareto distribution quantifying the heterogeneity in TCR immigration rates. The derived distribution for *π*_*α*_(*α*), given in Eq. 10 with *θ* = 0.28, however, is not large enough to generate a sampled clone abundance distribution 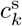 consistent with observations. In fact, the predicted distributions for reasonable sampling fractions *η* are nearly indistinguishable from those arising from a simple neutral model in which all clones have the same immigration rate. Conversely, proliferation heterogeneity within our model yields significantly different clone abundances that match those seen in experimental samples. Even under modest proliferation rate heterogeneity, larger clones become significantly more numerous at steady state since, although the number of TCR clones with large proliferation rates *r* may be small, such clones proliferate more rapidly contributing to higher clone counts at larger sizes. In particular, we found that the expected clone abundance is sensitive to the behavior of *π*_*r*_(*r ≈ R*), the proliferation distribution near the maximum proliferation rate *R*.

Our work leads to the conclusion that proliferation heterogeneity is the more likely mechanism driving the emergence of the power law distributions as observed in [13]. These results are consistent with the findings that peripheral selection leads to contraction of T cell diversity [49] and that naive T cells in humans are maintained by proliferation rather than thymic output [9]. Since we have only explored the effects of a uniform distribution for *π*_*r*_(*r*), further studies using more complex shapes of *π*(*α, r*) can be easily explored using our modeling framework. Different parameter values and rate distributions appropriate for mice, in which naive T cells are maintained by thymic output, should also be explored within our modeling framework.

## Acknowledgements

This work was supported by grants from the NIH through grant R01HL146552 and the NSF through grants DMS-1814364 (TC) and DMS-1814090 (MD). The authors also thank the Collaboratory in Institute for Quantitative and Computational Biosciences at UCLA for support to RD.

## S1: Mathematical Appendices

### A Neutral model

Here, we review the neutral model to provide insight into the properties of our heterogeneous BDI model. When there is no heterogeneity in either proliferation or immigration rates, 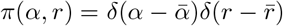. Upon inserting this expression for *π*(*α, r*) in Eqs. 7 and 8, we find that the clone abundance *c*_*k*_ follows a negative binomial distribution [10]:

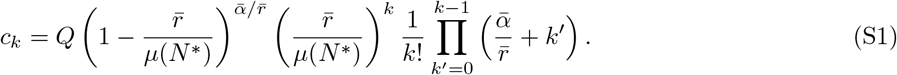

We can also express *c_k_/C*, the clone abundance distribution normalized by the mean richness *C* in the body as

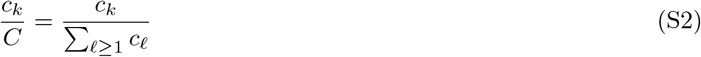

which is a negative binomial distribution of parameters 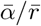 and 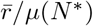. Using 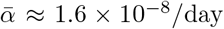, 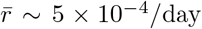, and *µ*(*N*\*) ≈ 6.4 × 10^*−*4^, we find 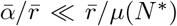. In this regime, *c_k_/C*, for *k* ≥ 1, can be approximated by a log-series distribution with parameter 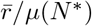. Fig. S2(a) shows that the exact solution for 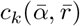 is indistinguishable from the log-series approximation.

To mathematically show that *c_k_/C* converges to a log-series distribution when 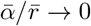, consider a random variable *X* that follows a negative binomial distribution of parameters 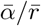 and 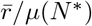

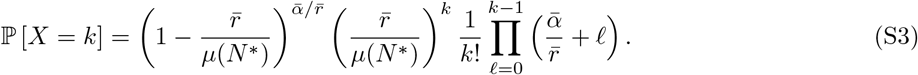

Note that the probability mass function of *X* is given by *c_k_/Q* as can be seen from Eq. S1, the clone abundance distribution for all possible *Q* clones, which includes *c*_0_, the number of all clones that are not represented in the organism. To find the clone abundance distribution *c_k_/C*, for all the *C* clones present in the organism, we must exclude the case *k* = 0 by marginalizing the distribution of *X* over all *X >* 0:

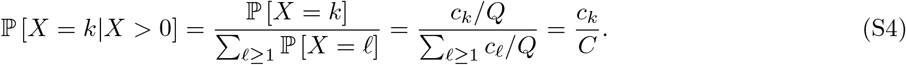

What remains is to show that the distribution of converges to a log-series distribution of parameter 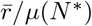 when 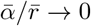. Consider the moment generating function of *X|X >* 0 given by

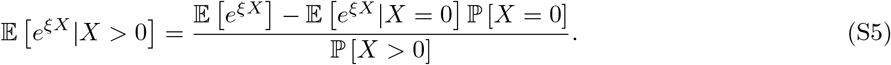

Since the moment generating function of a negative binomial distribution 𝔼[*e*^*ξX*^] is known, and since ℙ[*X >* 0] = 1 *−* ℙ[*X* = 0] (see Eq. S3), we can write

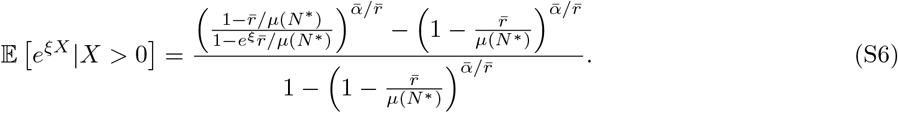

For any *x >* 0, the limit of 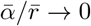, yields

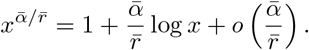

If we apply this result to Eq. S6 for 𝔼[*e^ξX^|X >* 0], we find

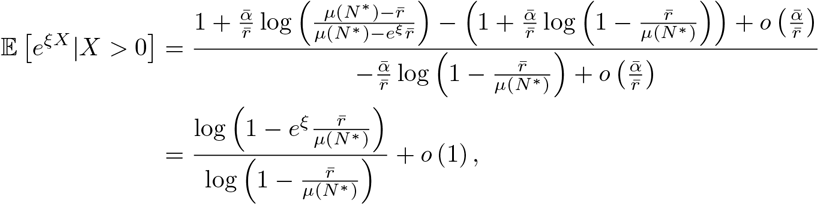

which we recognize as the moment generating function of a log series distribution of parameter 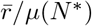. Thus, we finally have

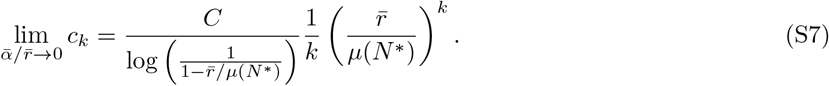

### B Heterogeneous immigration

Here, we provide technical details of the derivation of a Pareto distribution for the immigration rates *α*.

#### B.1 Determination of immigration rate distribution *π*_*α*_(*α*)

Marcou et al. [19] developed a computational tool, Inference and Generation of Repertoires (IGoR), which infers the probabilities of the recombination events that lead to the generation of a sequence. Using this tool, we generated 10^8^ different trials (for producing the human beta chain) and determined the frequency of occurrence of the corresponding sequence. Most of the sampled sequences arose only once, but others occurred twice or more in the sample. If *n*_*i*_ denotes the number of times each distinct sequence *i* arises in the 10^8^-trial simulation, its frequency is *f*_*i*_ = *n_i_/*10^8^. In Fig. S1 we plot the IGoR-generated frequencies *f*_*i*_ in descending order *f*_1_ *≥ f*_2_ *≥ … ≥ f*_*Q*_.

**Fig S1.**
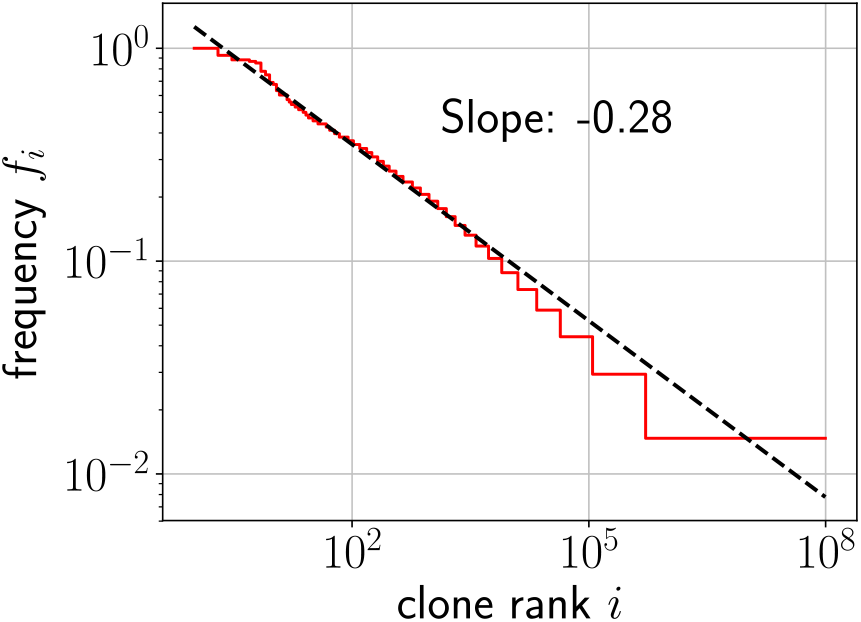
Clone rank versus frequency of sequences produced by IGoR (sampled for the human beta chain). Fitting this data to a Zipf distribution *f_i_ ∼ i*^*−θ*^ yields the best-fit parameter *θ ≈* 0.28.

By identifying the sequences generated by IGoR, we show that the distribution of thymic output rates follows a Zipf law (a discrete power-law distribution *f*_*i*_ ~ 1*/i*^*θ*^). This correspondence is not surprising since our procedure is analogous to determining word frequencies [45] from a given corpus of words in natural language, which is also known to follow a discrete power law. We can thus posit

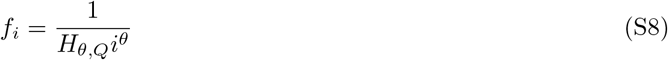

where 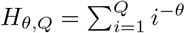 is a generalized harmonic number that serves as a normalizing constant. Using regression on the 10^8^ counts generated from IGoR, we determined that the slope of the Zipf distribution in our problem is *θ* ≈ 0.28.

Note that in principle we should probe a sufficient number of events to adequately represent the total number of distinct TCR sequences, on the order *Q* ~ 10^15^. However, since this is costly from a computational perspective, we generated 10^8^ recombination events (a “trial”) and assumed that the frequency-rank distribution would remain Zipf-like, with parameter *θ* ≈ 0.28, even under a higher number of trials. In the main text, we tested different values of *θ* in case this exponent is modified by choosing a larger number of trials.

Based on Zipf’s Law (Eq. S8), we reconstructed the statistics of immigration rates and *π*_*α*_(*α*) by associating *f*_*i*_ to the probability that each trial using IGoR generates a clone of sequence *i*. The total immigration rate 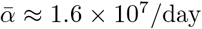 [37] represents the rate of the total TCR immigration. For each clone *i*, we can define its specific immigration rate *α*_*i*_ as

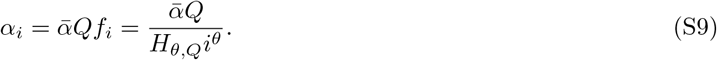

By definition, the immigration rates of all clones *α*_*i*_(1 ≤ *i ≤ Q*) are sampled from *π*_*α*_(*α*). Since the quantity *Q* is very large, one can assume that given bounds *a* and *b*,

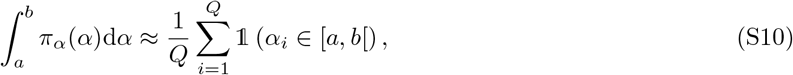

where 𝟙 (*α*_*i*_ ∈ [*a, b*[) is the indicator function, of unitary value if *α*_*i*_ falls in the [*a, b*[interval, and zero otherwise. From Eq. S8, we find the equivalent representation

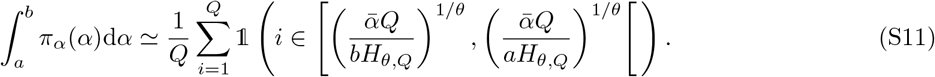

After using the first order approximation in the polynomial approximation for *b → a*, we find

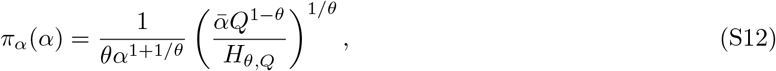

which is a Pareto distribution with parameters 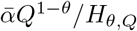 and 1*/θ*. Note that since *Q* is large, the exact computation of *H*_*θ,Q*_ is numerically challenging but a very accurate approximation can be obtained by considering its integral representation

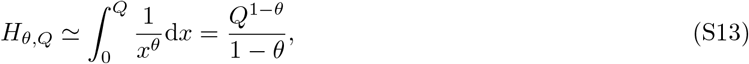

so that finally

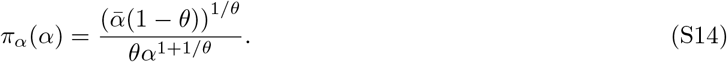

#### B.2 Shape of *c*_*k*_ in the heterogeneous immigration model

In the model with immigration heterogeneity, two regimes appear in the clone abundance *c*_*k*_ (see Figs. 2(b) and S2(b)). Small-sized clones are distributed similarly as in the neutral model; larger sized clones tend to follow a power-law. The interpretation of this behavior, provided in the main text, is that small-sized clones (population 1) are characterized by a low immigration rate 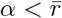, while larger-sized clones (population 2) are represented by a higher immigration rate 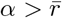. For clones in population 1, new cells arise in the clone mainly due to proliferation, while in population 2, they increase mainly due to immigration.

To verify this interpretation, we consider the clone count fraction *p*_*k*_ = *c*_*k*_/*C* for general *π*(*α, r*):

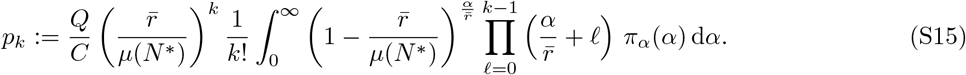

We separate the clone abundance distribution *p*_*k*_ into 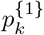, where immigration rates 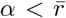, and 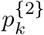 where immigration rates 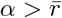 so that

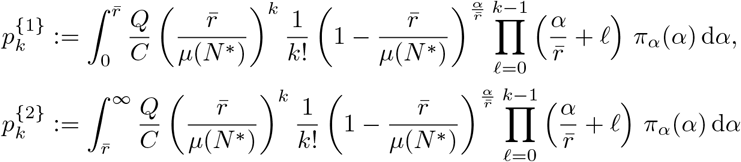

and 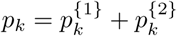(see Eq. S15). We approximate 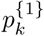 by focusing on the contribution to the integrand by terms with immigration rates 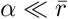, and similarly for 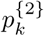 where we assume the greatest contribution arises from terms with 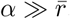.

To evaluate 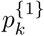, since 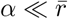 in the corresponding integrand, we can proceed in the same way as for the derivation of Eq. S7 in Section A to obtain a log-series distribution

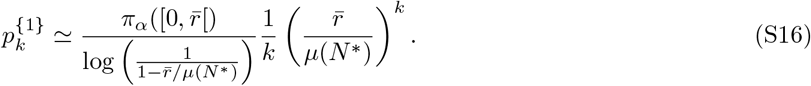

To evaluate 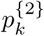, we consider a random variable 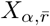 following a negative binomial distribution of parameters 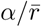 and 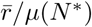 and note that 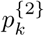 can be written as

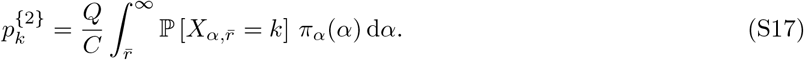

Since we have imposed 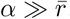, we consider the 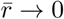 limit of 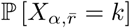, or equivalently the behavior of its moment generating distribution 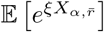 as 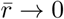. The latter is given by

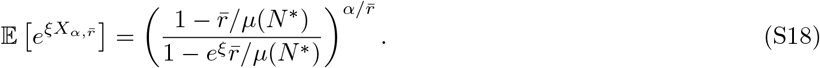

Under the assumption 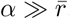, we take the 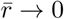 limit of Eq. S18 to find

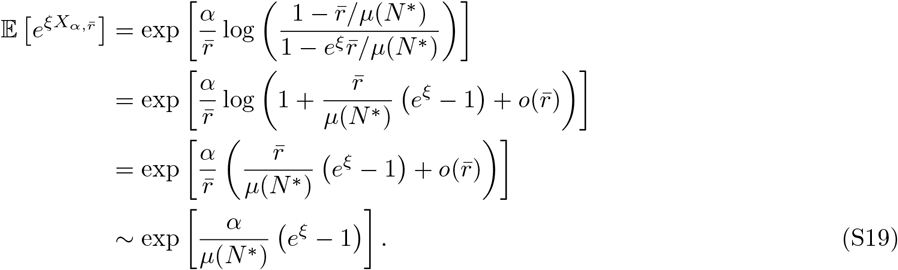

We now recognize the moment generating function of a Poisson distribution of parameter *α/µ*(*N*\*) so that

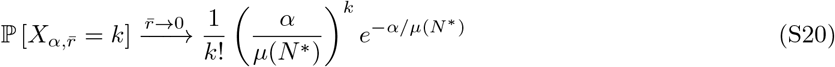

and

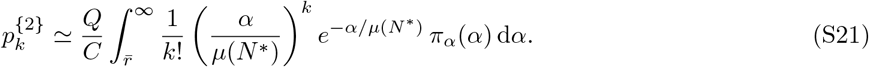

Thus, in this regime, clone abundance ratios obey a Poisson distribution of parameter *α/µ*(*N*\*). The approximations for 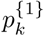 and 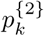 given by Eqs. S16 and S21 respectively are shown in Fig S2(b). Both approximations well represent the two sub-populations:

- Clones with a small cell number are mainly clones with an immigration rate 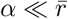 (population 1). When considering a clone in this regime that has at least one individual in the body, its distribution globally follows a log-normal distribution of parameter 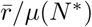. The creation of new cells in these clones is mainly driven by proliferation.
- Clones with a high cell number are mainly clones with an immigration rate 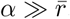 (population 2). When considering a clone in this regime with an immigration rate *α*, its distribution globally follows a Poisson distribution of parameter *α/µ*(*N*\*). The creation of new cells in these clones is mainly driven by thymic output.

**Fig S2.**
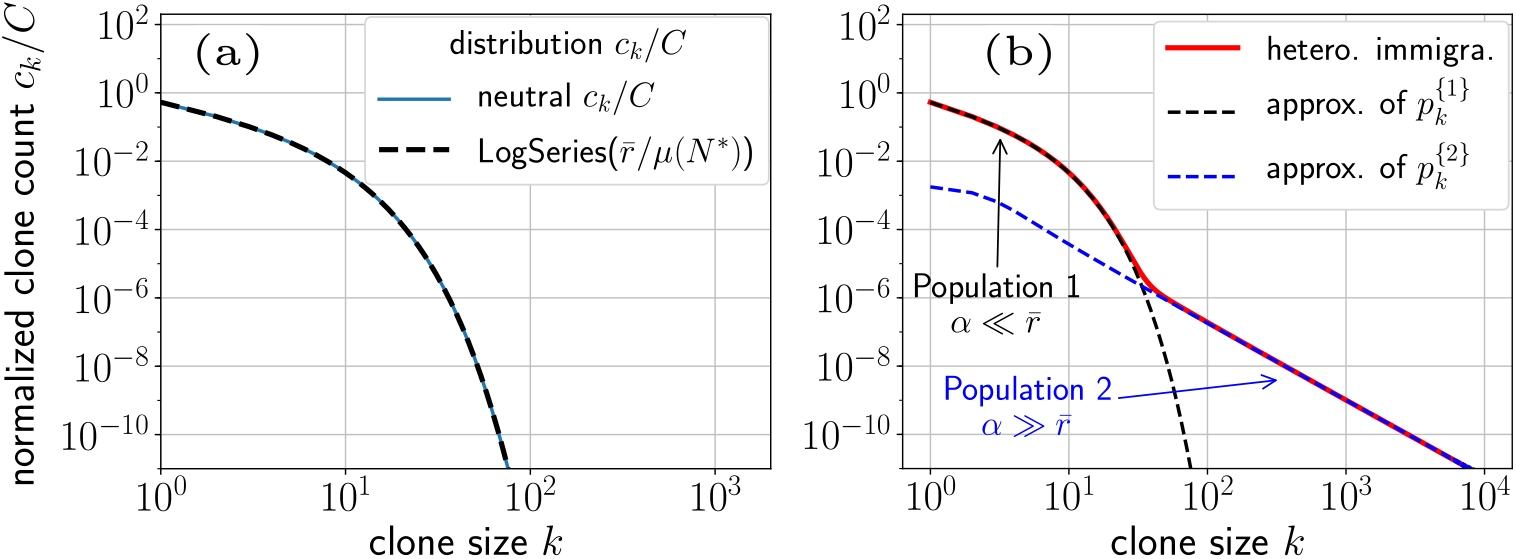
(a) Comparison of the clone abundance distribution (Eq. S7) with the log-series distribution in the neutral model. (b) Illustration of the two regimes in the clone abundance distribution of the heterogeneous immigration. The normalized clone abundance distribution *p*_*k*_ = *c_k_/C* is simply the normalization of *c*_*k*_ of Fig. 2(b) with *θ* = 0.8. The population *p*_*k*_ is subdivided in two sub-populations 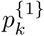 and 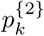 that represent clones with immigration rates *α* respectively lower or higher than 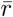 as described in Section B.2. The approximations of 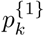 and 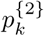(given by Eqs. S16 and S21), well represent the two regimes of *p*_*k*_.

**Fig S3.**
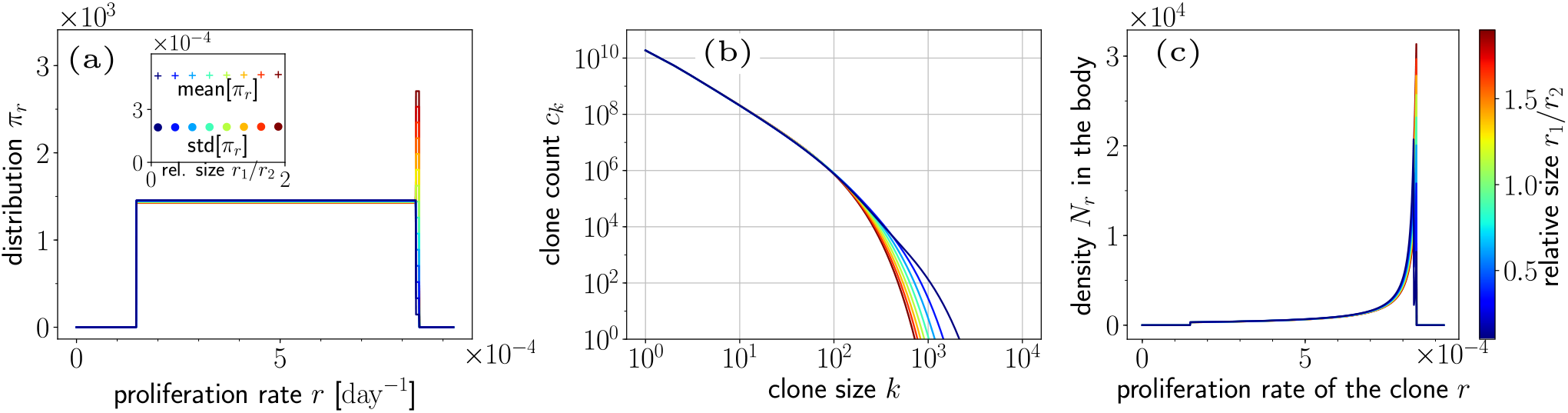
Effect of *π*_*r*_(*r ≈ R*) on the predicted mean clone count *c*_*k*_. (a) To study the effect of *π*_*r*_(*r*) when *r* is near its upper-bound *R*, we consider a series of almost-similar proliferation distributions *π*_*r*_(*r*) where only the local behavior for *r ≈ R* changes. Specifically, we allow *π*_*r*_(*r*) to take on two constant values: *π*_*r*_(*r*) = *r*_1_ for *r ≤* 0.99*R* and *π*_*r*_(*r*) = *r*_2_ for *R ≥ r ≥* 0.99*R*. By varying the ratio *r*_1_*/r*_2_, we change the local behavior at *r ≈ R* without significantly affecting the mean or the variance of *π*_*r*_(*r*) (see inset). (b) Decreasing *r*_1_*/r*_2_, increases the proportion of larger-size clones. (c) Spreading of proliferation rates *r* among the T cell population: *N*_*r*_ denotes the density of cells in the whole body with a proliferation rate between *r* and *r* + *dr*. Cells with higher *r* have a fitness advantage. Paradoxically, when *r*_1_*/r*_2_ decreases, the local competition between clones with proliferation rate *r ≈ R* decreases (the bluest curves are lower than the others around the upper bound *R*). Thus the fewer the number of clones with replication rate *r ≈ R* the less competition arises, leading to higher clone sizes *k* as observed in the (b).

**Fig S4.**
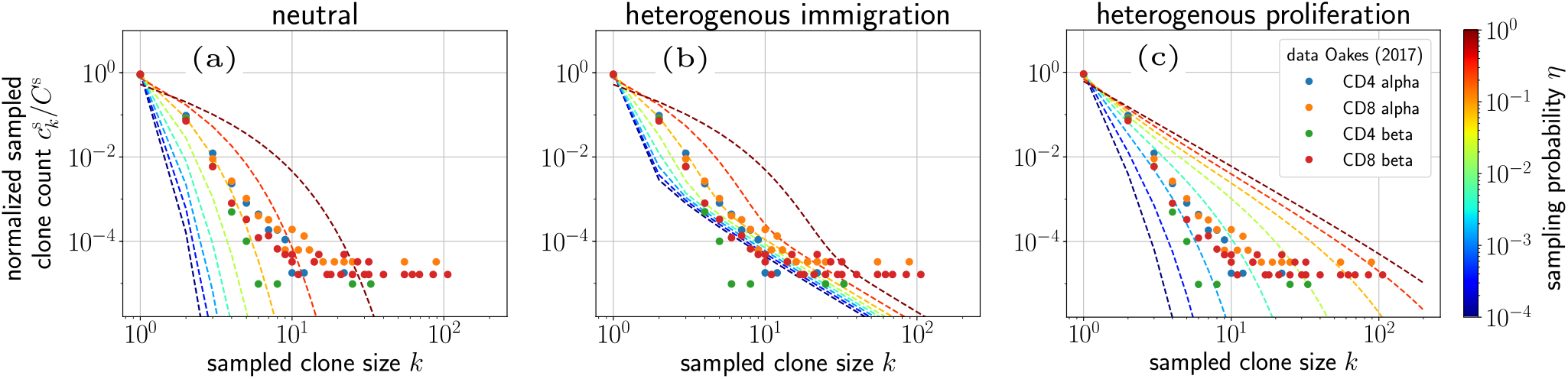
Effect of different sampling fractions *η* on predicted sampled clone abundances 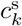. For all plots, the dots correspond to the experimental results of [13]. (a) Sampled clone abundances for the neutral model (without immigration or proliferation heterogeneity). (b) Sampled clone abundances for a model with high immigration rate heterogeneity corresponding to Fig. 2 for *θ* ≈ 1. The fit is poor even at this unrealistic value of *θ*. (c) Expected sampled clone counts for a high proliferation rate heterogeneity model (corresponding to the Fig. 3 for 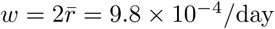).

**Fig S5.**
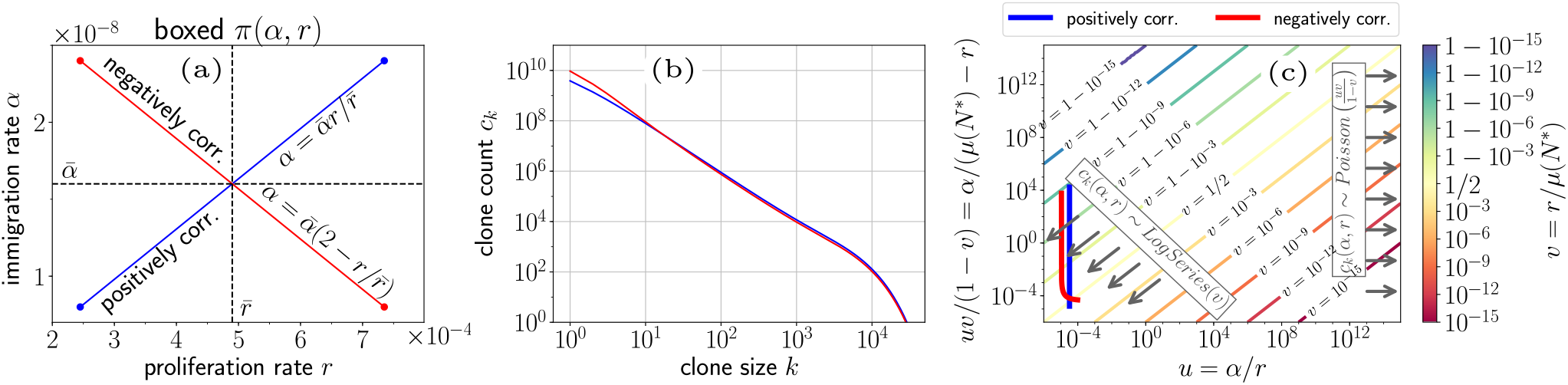
Positively and negatively correlated *π*(*α, r*). (a) For 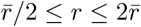, we consider *π*(*α, r*) distributions with positively and negatively correlated *α* and *r* (Eqs. 17). (b) Mean sampled clone counts corresponding to positively and negatively correlated *π*(*α, r*) show negligible differences. (c) “Line integrals” of the positively and negatively correlated distributions *π*(*α, r*) in the *uv/*(1 − *v*)- *u* diagram. Clones counts predicted by such *π*(*α, r*) follow log-series distributions, similar to those of a neutral model.

